# The balance between shear flow and extracellular matrix in ovarian cancer-on-chip

**DOI:** 10.1101/2023.07.20.549914

**Authors:** Changchong Chen, Alphonse Boché, Elliot Lopez, Juan Peng, Franck Carreiras, Marie-Claire Schanne-Klein, Yong Chen, Ambroise Lambert, Carole Aimé

## Abstract

Ovarian cancer is the most lethal gynecologic cancer in developed countries. Silent onset of the metastatic activity of ovarian tumor cells is factor for poor outcomes. In the tumor microenvironment, the extracellular matrix (ECM) and flow shear stress are known to play key roles in directing cell invasion. Hence, acute and tunable tools are critical to mimic scaffold and fluid for building clinically relevant *in vitro* models. We have built an ovarian tumor-on-chip where tunable ECM models are easily seeded with tumor spheroids and integrated within a microfluidic chip. This allows the investigation of the crosstalk between the characteristics of the ECM models and shear stress on the migratory behavior and cellular heterogeneity of ovarian tumor cells. We vary the composition of the ECM playing with type I and IV collagens and laminin, and control the shear stress in the chip. This work shows that in the shear stress regime of the peritoneal cavity, the ECM plays a major role in driving individual or collective modes of migration. In the presence of basement membrane proteins, migration is more collective that on type I collagen regardless of shear stress level. In addition, with increasing shear stress, individual cell migration was enhanced, while no significant impact on collective migration could be measured. This highlights our ability to discriminate relevant parameters for onset and shifts of cell behavior using our in vitro models. Furthermore, we described the ability to shift cells from an epithelial phenotype to a more mesenchymal phenotype, which could allow us to describe the role of these parameters during epithelial-to-mesenchymal (EMT) transition as a continuous process. Finally, we conclude that the ECM should hold a central position in *in vitro* cancer models, to understand cell response and develop platforms for therapeutic development.

## 1. Introduction

Ovarian cancer (OC) is the fifth most frequent cause of cancer death in women in developed countries [1]. This is attributed to the lack of visible and reliable symptoms during the onset of the disease, which results in patients being diagnosed at an advanced stage with increased probability of metastasis to distant organs. The biology of OCs differs from that of hematogenous metastasizing tumors because OC cells primarily disseminate within the peritoneal cavity. Most OCs result from the transformation of ovarian epithelial cells that grow into spheroids at a primary site and spread flowing through the pathological fluid that accumulates in the peritoneal cavity, called ascites [2]. In the course of this metastatic journey, tumor cells in their ascitic peritoneal environment are subjected to a low (less than a dyn.cm^-^²) isotropic shear stress resulting from walking and breathing movements [3]. Epithelial cells eventually undergo the epithelial-to-mesenchymal transition (EMT) and lose their epithelial characteristics for mesenchymal ones. This is associated with the weakening of their interactions with the basement membrane and the loss of cell to cell junction proteins like E-cadherin [4,5]. This works as a continuum, generating high cellular heterogeneity with intermediate cell states with varying expression of epithelial and mesenchymal markers. These hybrid states have been associated *in vitro* with improved metastatic properties including migration and invasion and increased cell survival in suspension. This also involves significant remodeling of the extracellular matrix (ECM), and the loss of basement membrane proteins, including collagen IV and laminin [6-8].

Advances in cancer research requires the ever improvement of *in vitro* models to reproduce the tumor microenvironment, which is known to impact cell proliferation, migration, invasion and apoptosis [9]. This includes (i) tridimensional cell constructs - spheroids, (ii) the circulating environment and (iii) the extracellular matrix (ECM). OC cells are likely to form spheroids, which may contribute to protect them from the circulating microenvironment [10] and to resist chemotherapies [11,12]. Many research efforts have been dedicated to the engineering of spheroid models using various strategies such as cell suspension culture, hanging drop, microwell arrays or microfluidics [13-19]. We have recently reported the microfabrication of non-adherent cages whose tunable dimensions enable tuning the spheroid size in a well-controlled and reproducible fashion [20]. In OC, the control over the circulating environment is particularly relevant, because OC cells benefit from the unique liquid microenvironment of the ascites exposing OC cells to shear stress. Microfluidic chips have mainly been developed to investigate the impact of shear stress on attachment, growth and motility of OC cells [21,22], the reorganization of the cytoskeleton under shear stress [23,24] and also the pathways involved in chemoresistance of non-adherent spheroids [25,26]. More recently, setups have been reported including vessels in multicellular constructs providing models to elucidate tumor-stromal cell interactions during intraperitoneal metastasis of OC [27-29]. Care must be also taken on the reconstruction of the surrounding ECM. This most often includes layer of fibronectin [21], growth factor reduced (GFR) Matrigel [27], and its mixture with collagen I [30,31], denuded amniotic membranes [24], fibrin hydrogels [29] and synthetic polymer coating [25,26].

Following previous works, we have built two ECM models made up of type I collagen, which is by far the major structural element of the ovarian stroma, eventually topped with layers of type IV collagen and laminin [32]. Collagen IV and laminin are two of the most abundant extracellular proteins in the mesothelium, which is extensively remodeled during OC. Collagen IV and laminin have been shown to be transiently lost in pre-malignant ovarian surface epithelium [6-8], which makes relevant to build models differing in their composition to mimic early and late stage of ovarian cancer. These ECM models exhibit stiffness in the range of the physiological, including pathological tissues, *i.e.* from few to tens of kPa [32]. This is made possible by the use of porous supports that maintain large areas of self-standing tissues, while allowing flow exchanges [33-35], and that can be easily inserted in a microfluidic.

By using only ECM proteins and different flow conditions, we aim to develop an ovarian cancer-on-chip and assess the crosstalk between ECM and shear stress in ovarian adenocarcinoma cell response. We show that migration strongly depends on the ECM, in the shear stress regime of the peritoneal cavity. Finally, in terms of morphology, the impact of the ECM was lowered under flow, and the shear stress appears to overcome the features of the ECM. These findings show the unique ability of our model to uncouple biochemical and physical cues and analyze their impact on the migratory behavior and morphology of cells.

## 2. Materials and methods

### 2.1. Fabrication of the porous supports

Porous supports similar to the ones that are commercially available from the Mesobiotech® company were fabricated by photolithography and soft-lithography in the lab as previously reported (see Supporting Information) [32]. The width of the honeycombs was set at 400 µm, with a frame width of 50 µm and a thickness of 50 µm.

### 2.2. ECM fabrication

Type I collagen was extracted and purified from rat tail tendons as previously described, except that we used 3 mM hydrochloric acid instead of 500 mM acetic acid [36,37]. Collagen purity was assessed by electrophoresis and its concentration estimated by hydroxyproline titration [38]. All other chemicals were purchased and used as received. Water was purified with a Direct-Q system (Millipore Co.). Two ECM models were prepared. The first one is an artificial connective tissue (ACT) from type I collagen only. A PBS solution at pH 9 was prepared by slowly adding 1M NaOH to 1X PBS (pH = 7.4) to induce collagen I fibrillogenesis from acidic collagen I solution (1 mg/mL in PBS). In the second model, called artificial basement membrane (ABM), type I collagen (1 mg/mL in PBS), type IV collagen (0.5 mg/mL in PBS, from human placenta) and laminin (10 μg/mL in dionized (DI) water, from Engelbreth-Holm-Swarm murine sarcoma basement membrane) were drop cast by pouring 20 µL of the solution on the patch and dried in air at room temperature in between each step of drop casting in the layer-by-layer process (Figure 1A). Finally, the ECM was washed with DI water, dry in air at room temperature and could be stored in the fridge at 4℃.

**Figure 1.**
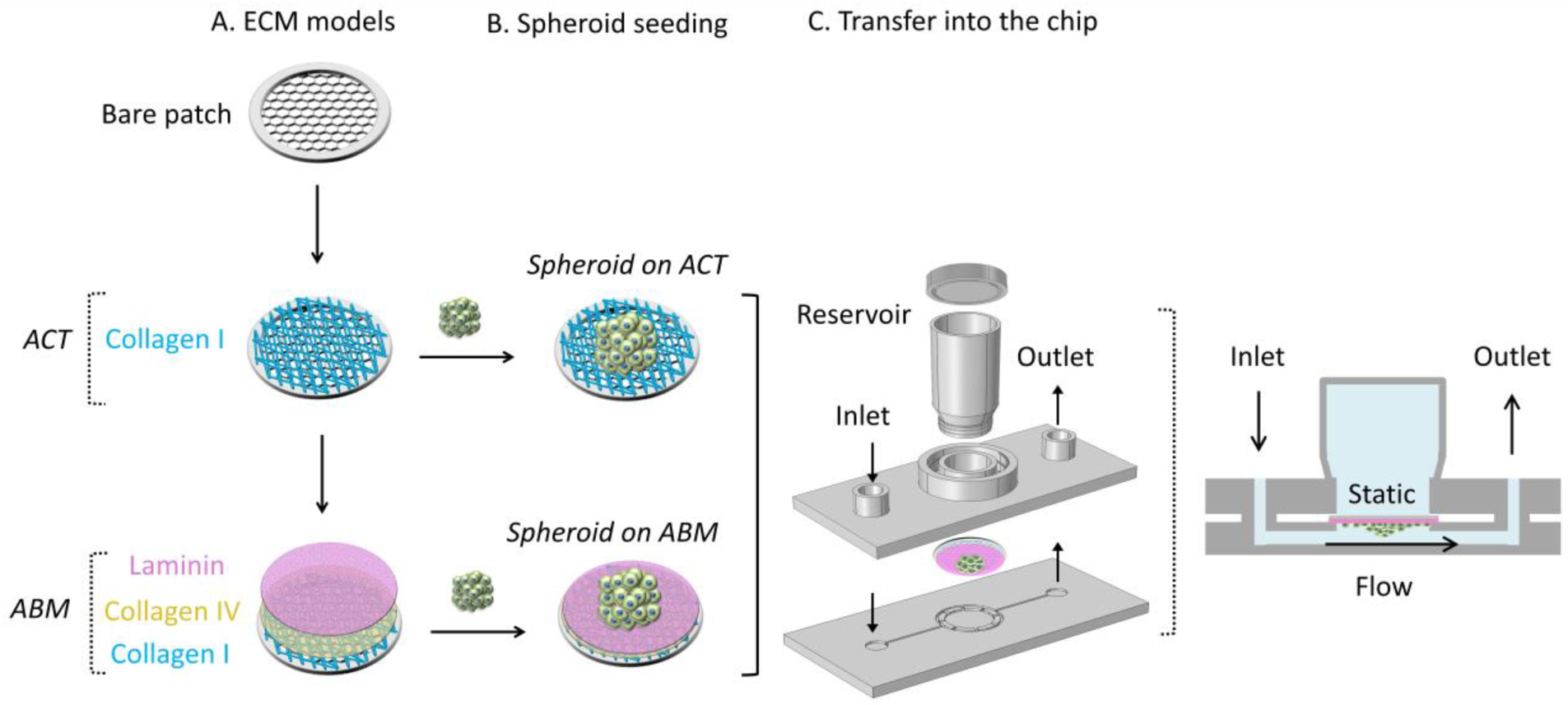
Scheme of the OC-chip. (A) ECM models: artificial connective tissues (ACT) and artificial basement membrane (ABM), (B) spheroids seeding, and (C) integration of the seeded ECM into the microfluidic chip.

### 2.3. Cell Culture

Human ovarian adenocarcinoma SKOV-3 cells (ATCC1, HTB77™) were purchased from ATCC (American Type Culture Collection, Manassas, VA). Cells were cultured in RPMI-1640 glutaMAX containing 0.07 % (v/v) sodium bicarbonate supplemented with 10% fetal bovine serum and 1% (v/v) penicillin streptomycin (all reagents were purchased from Thermo Fisher Scientific). Cells were cultured in T25 cell culture flasks in a humidified air atmosphere with 5% CO_2_ at 37°C.

### 2.4. Spheroid growth

Ovarian tumor spheroid growth was performed with a high control over spheroid size, as recently described [20]. Briefly, gelatin nanofibers were electrospun on caged-patch having a wall thickness of 200 µm. Then, agarose was coated on the gelatin fibers. After maintaining the spherical patch in the bottom of a 12-well plate using a PDMS ring, SKOV3 cells (4×10^5^ cells/patch) were deposited on the caged-patch. After 3 days culture, uniform spheroids were obtained with a diameter of *ca.* 200 μm. Ovarian tumor cells in spheroids were stained with epithelial (EpCAM and E-cadherin) and mesenchymal (Vimentin) markers (Figure S1-S2).

### 2.5. Seeding ovarian tumor spheroids and integration in the microfluidic chip

Cell migration experiments were carried out on ACT and ABM and in static and flow conditions. First, all ECM models were sterilized by immersion in 70% ethanol for 5 min before washing in sterilized PBS 3 times for 5 min each time. The ACT/ABM supports were maintained at the bottom of the dish with a PDMS ring. Two to four SKOV3 tumor spheroids were selected out using a micropipette and seeded one by one (Figure 1B). The tumor spheroids were incubated on the ECM models for 2 hours at 37℃ and 5% CO_2_.

The seeded ECM was then transferred in a commercial microfluidic device (Mesobiotech, France) composed of two plastic plates and a circular culture chamber (∅ 10 mm, h 1.5 mm) at the center of the chip (Figure 1C). The chip is closed with magnetic bounding and a mechanical clamper with four hand screws to prevent leakage for cell culture under flow for two days. The upper plate was designed to have two Luer connectors (one inlet and one outlet) and a central reservoir. The inlet channel is divided into six smaller channels to spill the flow into the central cavity while reducing the shear stress. The distribution of the shear rate (ν in s^-1^) in the central cavity was simulated by computer fluid dynamics modeling for two different flow rates and calculated to be ν ɛ (0.2-0.4 s^-1^) at 20 µL.min^-1^ and ν ɛ (1-2 s^-1^) at 100 µL.min^-1^ (see Supporting Information). Accordingly,

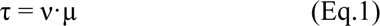

where τ is the shear stress and µ is the dynamic viscosity of culture medium. Thus, τ ɛ (2.4×10^-3^-4.8×10^-3^ dyn.cm^-2^) at 20 µL.min^-1^ and τ ɛ (1.2×10^-2^-2.4×10^-2^ dyn.cm^-2^) at 100 µL.min^-1^, respectively, with a dynamic viscosity of 1.2×10^-2^ dyn.s.cm^-2^. For the given channel with a section area in the central chamber of ∼0.2×10 mm², a linear fluid velocity of ∼0.2 and 0.8 mm.s^-1^ can be obtained at flow rate of 20 and 100 µL.min^-1^ respectively, which also give a shear rate in the order of the simulated values.

### 2.6. SEM imaging

For ECM characterization, samples were dried at 60°C overnight, then coated with gold for 60 s using an Emitech K675X Sputter Coater System with a sputtering current of 50 mA before imaging. Samples were fixed on conductive-tapes for imaging with a TM3030 Tabletop Microscope (Hitachi High-Technologies Corporation, Japan) working at an acceleration voltage of 15 kV.

### 2.7. Second harmonic generation / 2-photon excited fluorescence

We used a custom-built laser-scanning multiphoton microscope and recorded second harmonic generation (SHG) and 2-photon excited fluorescence (2PEF) images in parallel as previously described.^39^ Excitation was provided by a femtosecond titanium–sapphire laser (Mai-Tai, Spectra-Physics) tuned to 860 nm, scanned in the XY directions using galvanometric mirrors and focused using a 25× with 1.05 NA objective lens (XLPLN25X-WMP2, Olympus), with a resolution of 0.35 μm (lateral) × 1.2 μm (axial) and a Z-step of 0.5 µm for the acquisition of Z-stack images. We used circular polarization in order to image all structures independently of their orientation in the image plane, using 100 kHz acquisition rate, 420×420 nm² pixel size and a typical power of 22 mW.

For the characterization of the basement membrane by 2PEF, ECM models were stained with Alexa Fluor 488^®^-conjugated collagen IV monoclonal antibody (eBioscience™, 53-9871-82) incubated overnight at 4°C or 2 h at room temperature at a concentration of 10 μg.mL^-1^, together with laminin polyclonal primary antibody (Thermo Fisher Scientific, PA1-16730) overnight at 4°C or 2 h at room temperature at a concentration of 1 μg.ml^-1^, followed by 1 to 2 h incubation at room temperature with a goat–anti rabbit-Alexa Fluor 610 secondary antibody at a dilution of 1/250.

### 2.8. Immunofluorescence and confocal image

After 2 days of culture, cells were fixed in 4% paraformaldehyde (PFA) in PBS for 10 minutes, rinsed three times with PBS for 5 min each time. The cells were permeabilized with 0.5% Triton X-100 in PBS for 10 minutes, washed again 3 times in PBS and saturated with PBS containing 3% BSA (with 0.1% Tween 20 and 0.1% sodium azide) for 30 min and washed again. Cells were then incubated overnight at 4°C with Alexa Fluor 543-conjugated vimentin antibody (ab202504, Abcam®) at a 1/1000 dilution and washed 3 times with PBS for 5 min each time to remove unbound vimentin conjugate. After washing, the actin cytoskeleton was stained with a 50 mg/mL Alexa Fluor 488 Phalloidin in PBS (containing 1% DMSO from the original stock solution, Abcam®) for 40 min at room temperature in a dark chamber. Cells were washed 3 times with PBS for 5 min each time to remove unbound phalloidin conjugate. Nuclear DNA was then stained with DAPI (4,6-diamidino-2-phenylindole dihydrochloride, Molecular Probe®) for 15 min at room temperature and washed again. Immunofluorescent labelling was observed with a confocal microscope (LSM710, Zeiss) equipped with 405 (DAPI), 488 (phalloidin), and 543 (anti-vimentin antibody) nm lasers and with LSM ZEN 2009 software. We used 1 μm z-stack intervals and sequential scanning for each wavelength.

### 2.9. Image processing and statistical analysis

The migratory behavior (migrating distance, number of migrating cells and core area) was analyzed using Image J [40]. The staining intensity of vimentin in the core and leading areas were analyzed using imageJ by measuring the ratio of the integrated fluorescence intensity of a region of interest to the area of this region. More than 4000 cells were analyzed. Every group of experiments has been run in triplicate and at least 9 tumor spheroids were statistically analyzed for each condition using the Two-Sample t-Test. Finally, the morphology of cells and nuclei were analyzed by principal component analysis (PCA) using the CellTool software [41,42]. PCA allows the determination of the standard deviation (s.d.) that represents the squareroot of the total variance calculated from all the measured morphological features, and hence quantifies the cellular heterogeneity.

## 3. Results

### 3.1. Impact of the ECM model on the migration of ovarian tumor spheroids

To construct the ovarian tumor-on-chip, porous supports with a diameter of 1.3 cm and a honeycomb microframe were fabricated that can be easily integrated into the circular chamber of our microfluidic device (Figure 1). The high porosity of the frame (400 µm-width honeycombs) ensures fluid exchanges and an important area of self-supporting tissue with limited mechanical contribution of the support (Figure 2). Two ECM models were selected to investigate the impact of key proteins in OC environment (Figure 1A). The first one, referred to as artificial connective tissues (ACT) is made up of a fibrillar type I collagen network. This reproduces the late-stage of OC, characterized by the loss of key proteins from the basement membrane. The second one is made up of a fibrillar network of type I collagen topped with type IV collagen and laminin to figure an artificial basement membrane (ABM). SEM imaging of the ACT shows the self-assembly of a dense layer of type I collagen fibers on the honeycomb microframe (Figure 2A1). For imaging the ECM in hydrated conditions, we used second harmonic generation (SHG), which is highly specific to the organization of collagen into densely packed triple helices. Figure 2A2 shows that the collagen I layer is homogeneous, with a thickness of a couple of microns. After adding successively type IV collagen and laminin, the layer thickness and aspect did not change under SEM (Figure 2B1). Combining SHG with 2-photon excited fluorescence (2PEF) allows to specifically identify the presence of each protein: a top layer of laminin, with type IV collagen embedded within the ABM as observed on the different isolated channels (Figure S4).

**Figure 2.**
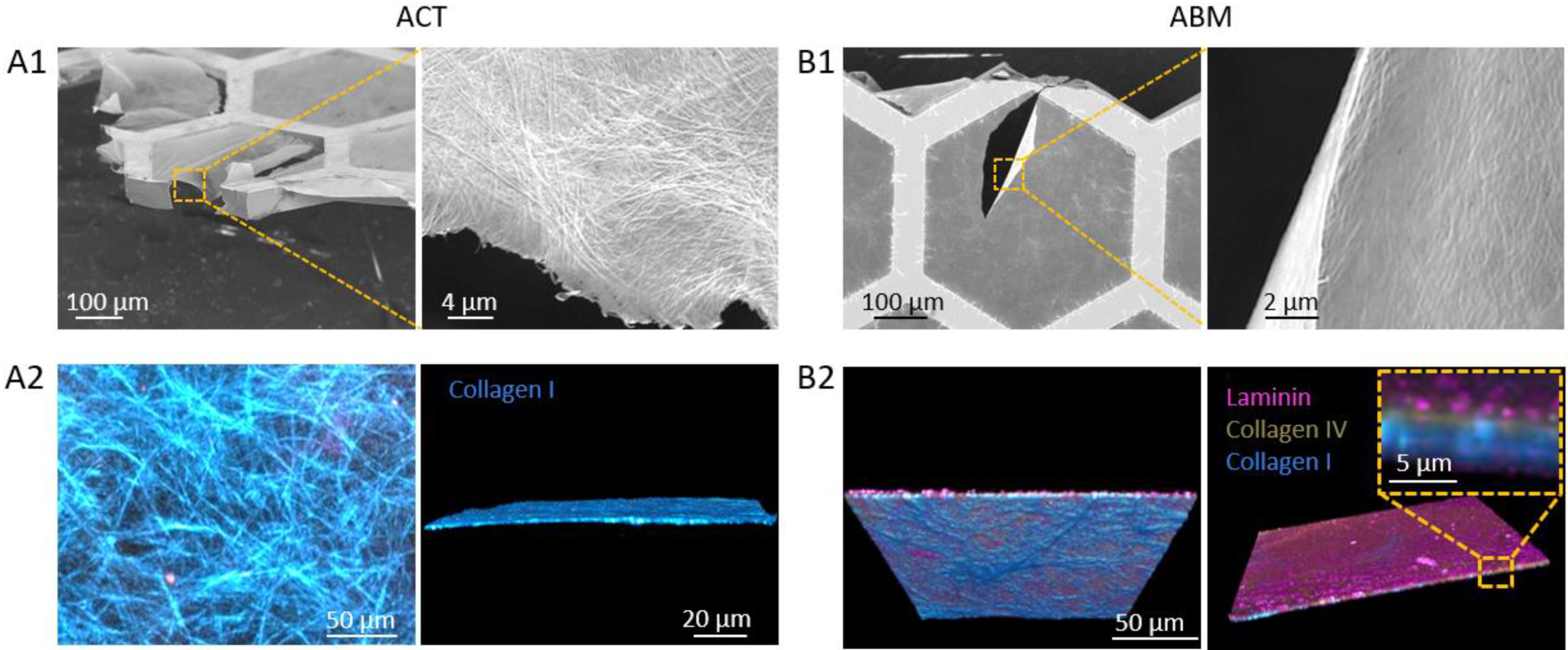
(A1,B1) SEM and (A2,B2) SHG/2-PEF images of (A) ACT and (B) ABM on honeycomb frame supports.

Ovarian tumor spheroids (OTS) with a uniform diameter of about 200 µm [20] were seeded on ACT and ABM (Figure 3), and the impact of the ECM was evaluated in static conditions after immunofluorescence (IF) imaging. To quantify the migratory behavior, the mean migration distance, the number of migrating cells and core area were measured. We define the mean migration distance as the distance from the core to the front of the spheroid. Since the honeycomb frame has six-fold symmetry, we averaged the migration distance over 8 positions to account for the unsupported ECM and the frame (see yellow arrows in Figure 3A). The number of migrating cells was defined by the number of individual cells out of the spheroid core. On ACT models, the mean migration distance was found to be significantly higher than on ABM, and associated to a significantly lower number of migrating cells (Figure 3D,E). This signs for cells having a higher motility and more individual migratory behavior on ACT compared to the collective migratory behavior on ABM in static conditions and show that OC cells are strongly affected by the ECM. This is well-illustrated in Figure 3B-C, where a distinct spheroid core with isolated surrounding cells can be observed on ACT in contrary to the large growing core observed on ABM. Indeed, significantly smaller core area of spheroids were measured on ACT compared to ABM (Figure 3F). Zoom-in images show the phenotypic variations of cells together with the variation of vimentin staining (in red). Vimentin is a cytoskeleton protein upregulated during cell transitioning events and commonly used as a mesenchymal marker [43-46]. E-cadherin and EpCAM were also systematically investigated as epithelial markers, staining cell-cell junctions (Figure S5). Typically, on both ECMs, cells in the core of spheroids were found to be round with a low expression of vimentin and becomes elongated from the core to the front of the spheroid, with increased expression of vimentin (Figure 3G). This expression pattern shows the prevalence of epithelial-type cells in the core of the OTS and evolving towards mesenchymal phenotypes at the migration front, which was enhanced for ABM compared to ACT (Figure 3G).

**Figure 3.**
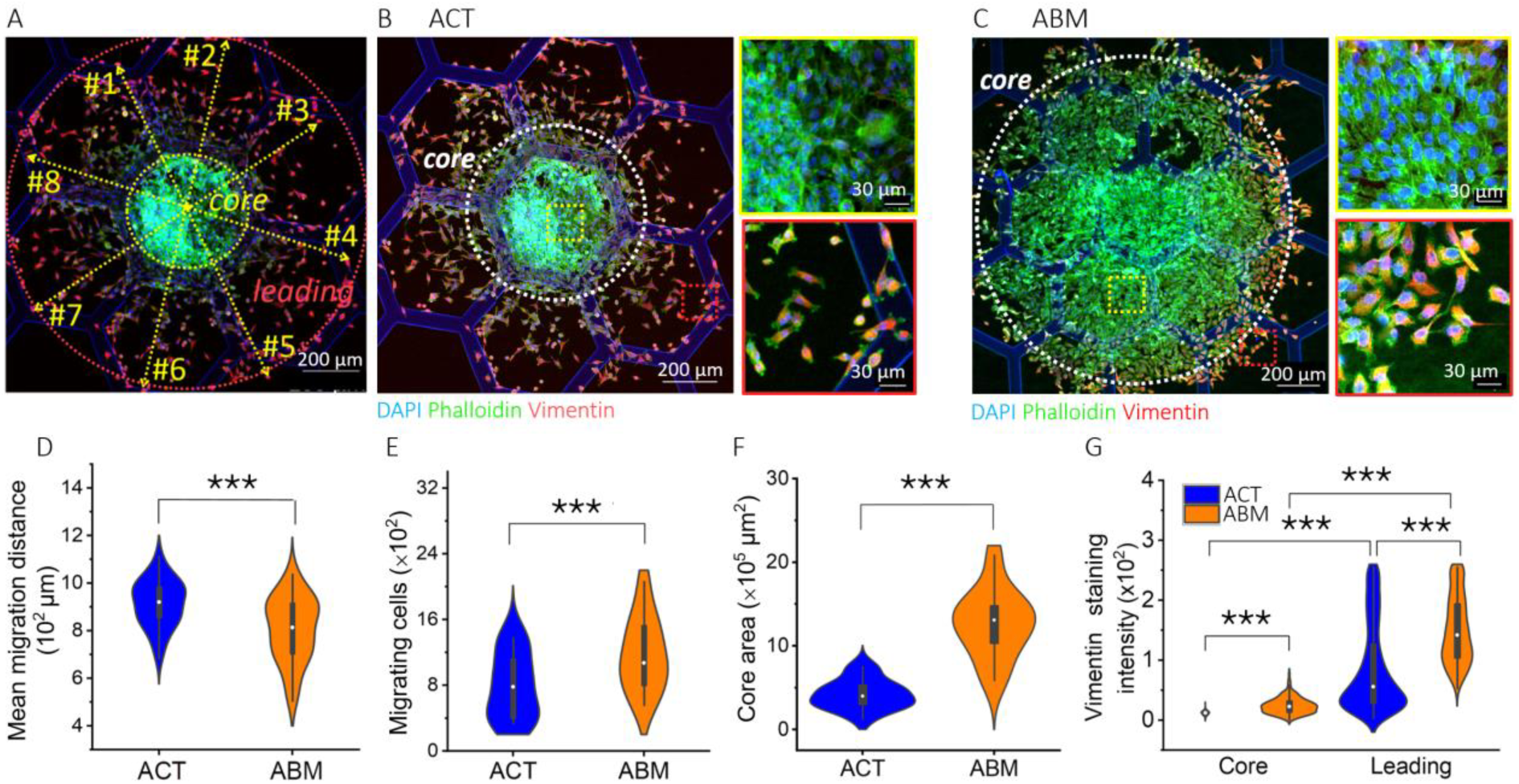
(A) Scheme of the measurement of the mean migration distance, number of migrating cells and core area. (B,C) IF imaging of SKOV-3 cells after 48 hours culture on ACT and ABM respectively. Cells were stained for vimentin (red), actin (green), and nuclei (DAPI, blue). (D-F) Plots of the mean migration distance, number of migrating cells and core area on ACT and ABM models in static conditions. (G) Vimentin staining intensity as a function of cell positioning in the spheroid (***p < 0.001).

We further analyzed the morphology of isolated cells (in the leading region) and their nuclei using principal component analysis (PCA) (Figure 4) [41,42]. PCA provides information about the number of geometrical descriptors -*shape modes*-that account for the morphological heterogeneity of a population of cells/nuclei. In addition, for each descriptor it gives the distribution of values associated with standard deviations. Cell morphology, based on the staining of the actin cytoskeleton, reveals four shape modes on ACT and five on ABM, accounting for a higher heterogeneity on ABM (Figure 4A). The most important geometrical descriptor for both cell populations is the elongation of cell shape, being more elongated for cells on ACT than on ABM. Nuclei were also found to be more elongated on ACT than on ABM in static conditions, with smaller size (Figure 4B).

**Figure 4.**
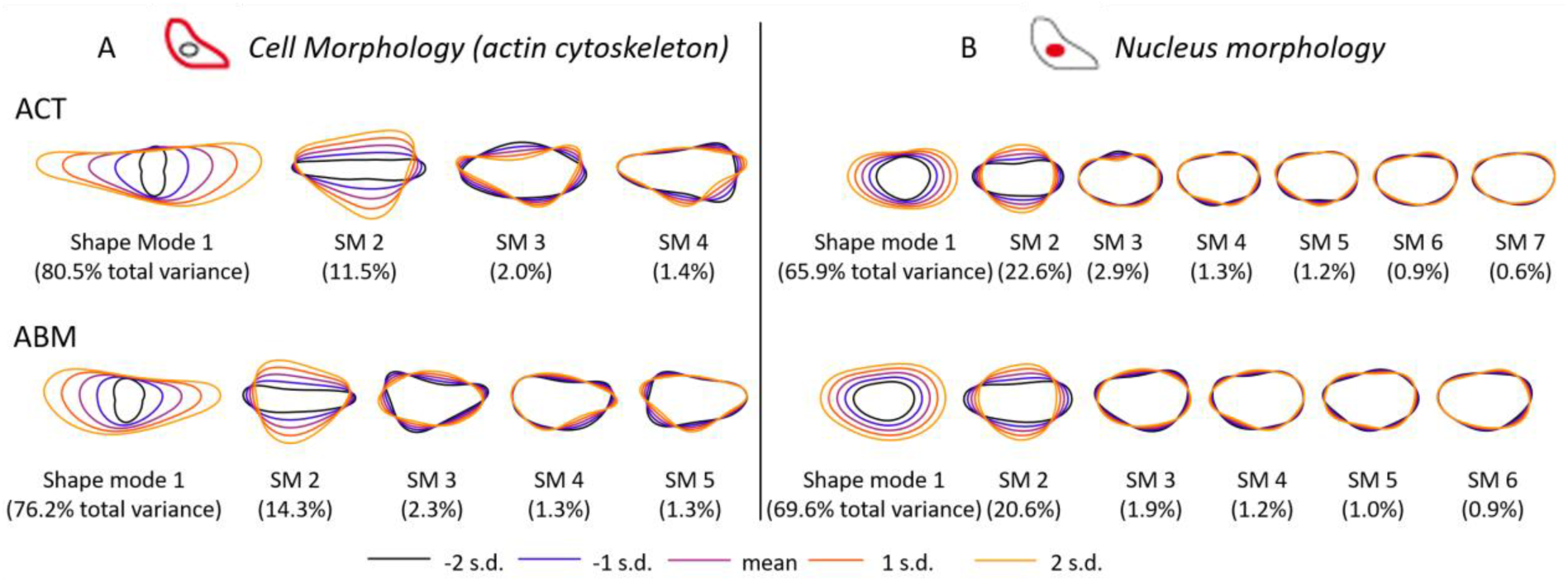
PCA of nucleus and cell morphology 48h after seeding on the different ECMs in static conditions: (A) morphological variance of cell morphology based on phalloidin staining of the actin cytoskeleton and (B) morphological variance of nucleus morphology based on DAPI staining. The color code is common to all plots. s.d. is for standard deviation, as determined by PCA for the quantification of the variance in the measured morphological features.

### 3.2. Effect of fluid shear stress on ovarian tumor cells cultured on ACT

After the identification of ECM impact on SKOV-3 spheroid behavior, we added up flow to make cells experience the fluid shear stress of the peritoneal cavity. Compared to the conventional approach relying on cell culture with an already integrated membrane, we use a “seeding first and insert after” approach, where the culture support engineered with the ECM and seeded with OTS is inserted in the central chamber of the chip (Figure 1). This also allows easy removal of the culture support for downstream IF observations. Two perfusion speeds were selected to match the physiological shear stress measured in dyn/cm² for biological applications and which depends on cavity and vessel types, being less than a dyn.cm^-^² in the peritoneal cavity. Perfusion speed of 20 and 100 µL.min^-1^ were calculated to give shear stress values in the range of 10^-3^ and 10^-2^ dyn.cm^-2^ respectively, which correspond to values of shear on cells of about 0.2 and 0.8 mm.s^-1^. In both conditions, the individual migration mode observed under static conditions was confirmed. Spheroids show a small and dense core with isolated cells migrating towards the front (Figure 5A,B). The mean migration distance and number of migrating cells were found to decrease with increasing shear stress, together with a smaller area of the spheroid core (Figure 5C-E). Regarding the intensity of vimentin staining, an important increase was again observed from the spheroid cores to the front, whatever the shear stress (Figure 5F). This signs for the acquisition of mesenchymal markers in the course of migration. No increase in vimentin staining could be measured in the core upon increasing the shear stress. However, the intensity of vimentin staining in the leading area of the spheroid was found to be significantly enhanced upon increasing the shear stress (Figure 5F). This fosters the individual migration mode, where the number of migrating cells decreases with enhanced mesenchymal properties as highlighted by the intensity of vimentin.

**Figure 5.**
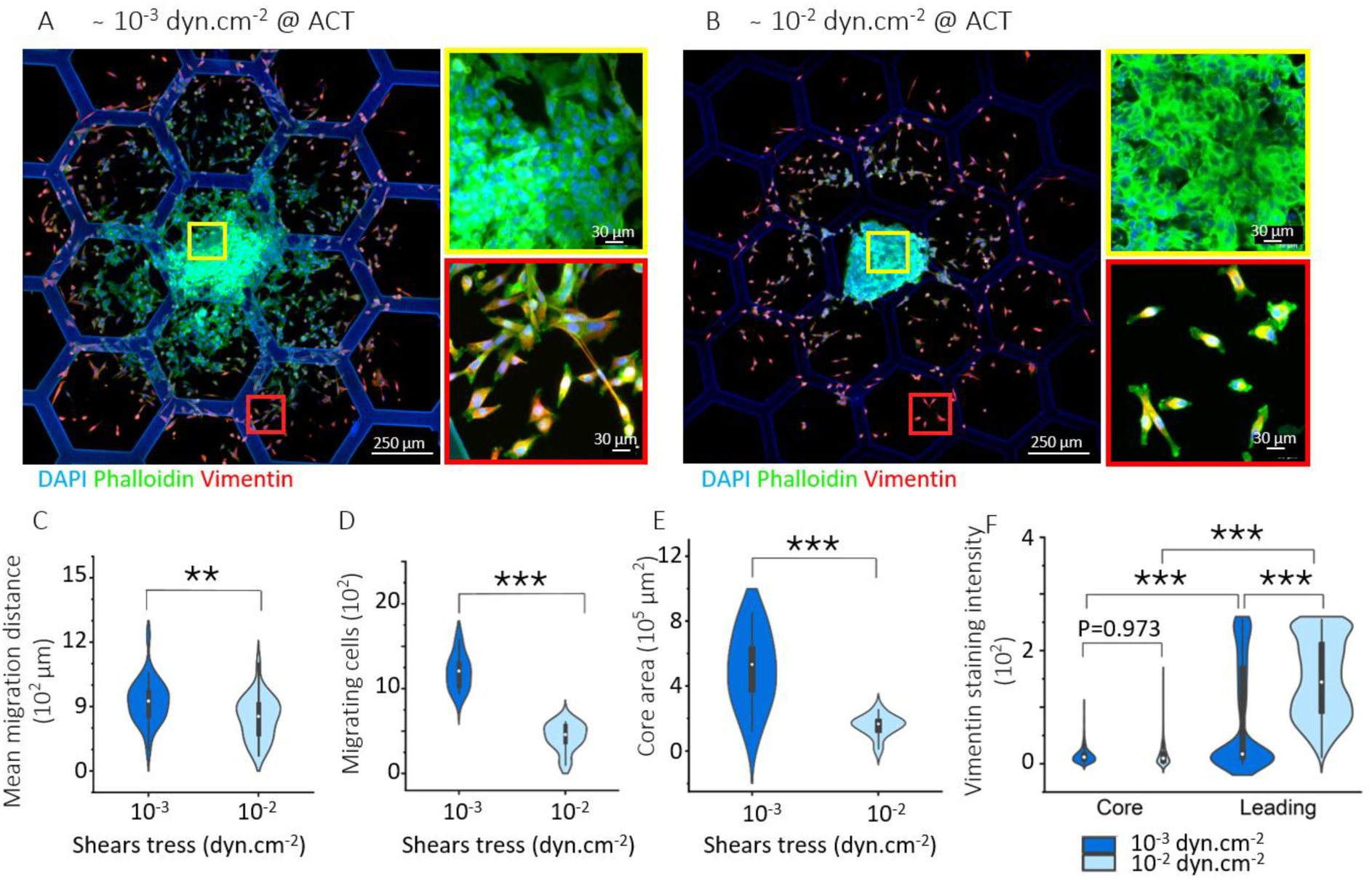
Effects of flow shear stress on migration of SKOV-3 cells on ACT. (A,B) IF imaging of SKOV-3 cells after 48 hours culture on ACT. Cells were stained for vimentin (red), actin (green), and nuclei (DAPI, blue). (C-E) Plots of the mean migration distance, number of migrating cells and core area on ACT as a function of shear stress. (F) Vimentin staining intensity as a function of cell positioning in the spheroid and shear stress (**p <0.01; ***p < 0.001).

The heterogeneities measured in terms of cell and nucleus morphology were found to be more important at high shear stress. In the shear stress range of 10^-3^ dyn.cm^-2^, three and five shape modes describe the population of cells and nuclei respectively, compared to four and six shape modes in the 10^-2^ dyn.cm^-2^ range (Figure 6). Moreover, both cell and nuclei were found to decrease in size, while being more elongated when increasing the shear stress from 10^-3^ to 10^-2^ dyn.cm^-2^.

**Figure 6.**
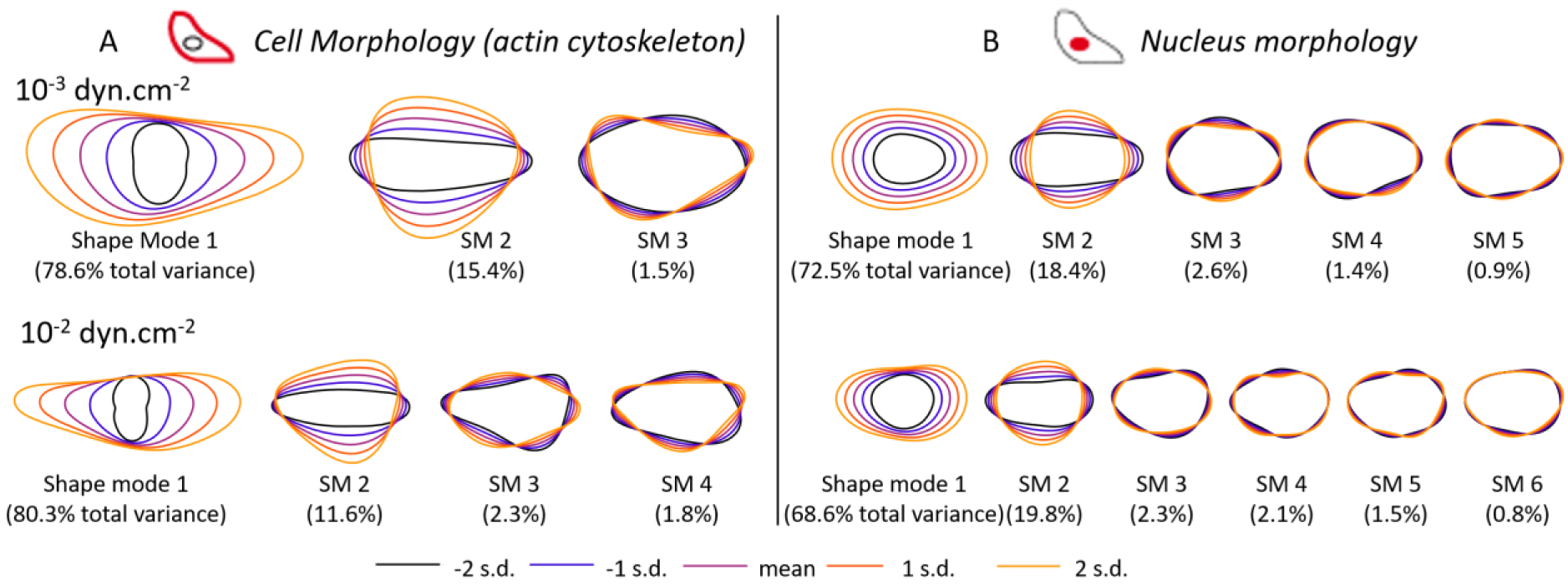
PCA of nucleus and cell morphology 48h after seeding on ACT as a function of flow shear stress: (A) morphological variance of cell morphology based on phalloidin staining of the actin cytoskeleton and (B) morphological variance of nucleus morphology based on DAPI staining. The color code is common to all plots. s.d. is for standard deviation, as determined by PCA for the quantification of the variance in the measured morphological features.

### 3.3. Effect of fluid shear stress on ovarian tumor cells cultured on ABM

SKOV-3 spheroids were finally seeded on ABM and cultured under flow at 10^-3^ to 10^-2^ dyn.cm^-2^ for two days. The collective migration observed under static conditions was confirmed under flow, with large spheroids surrounded by few isolated cells (Figure 7A,B). While the mean migration distance was found to slightly increase with the shear stress, the number of migrating cells and core area were not significantly affected indicating a limited impact of the shear stress on the migration behavior of OC cells cultured on ABM (Figure 7C-E). The staining intensity of vimentin was found to again increase from the core to the leading area of the spheroid but with a rather limited increase in the leading region with the shear stress compared to the increased observed on the ACT scaffold (Figure 7F).

**Figure 7.**
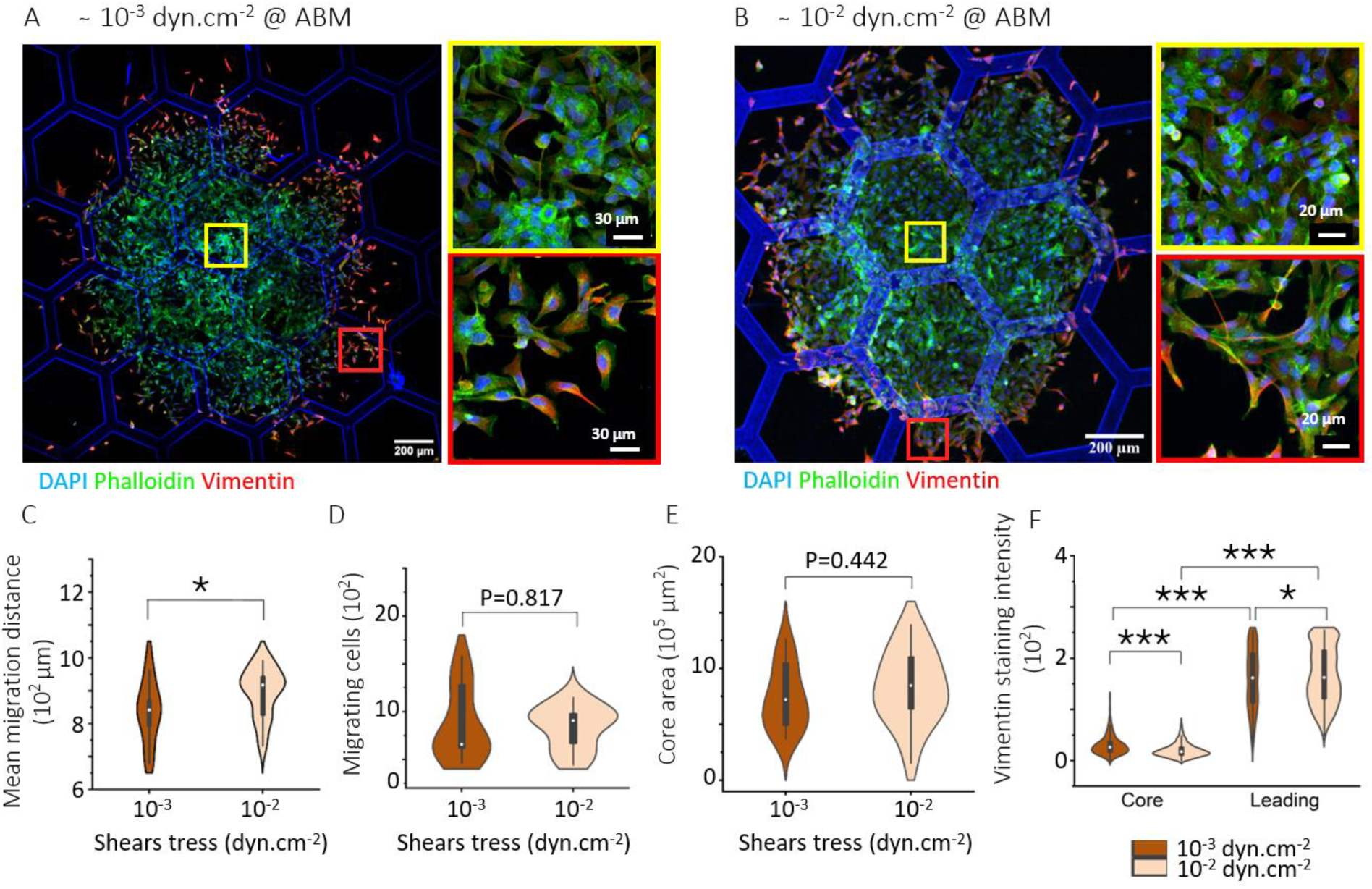
Effects of flow shear stress on migration of SKOV-3 cells on ABM. (A,B) IF imaging of SKOV-3 cells after 48 hours culture on ABM. Cells were stained for vimentin (red), actin (green), and nuclei (DAPI, blue). (C,E) Plots of the mean migration distance, number of migrating cells and core area on ABM as a function of shear stress. (F) Vimentin staining intensity as a function of cell positioning in the spheroid and shear stress (*p <0.05; ***p < 0.001).

Finally, the heterogeneity of cell and nucleus morphology upon increasing shear stress was similar with four shape modes describing the heterogeneity of cell morphology whatever the shear stress and four and five shape modes accounting for the nucleus heterogeneity at 10^-3^ to 10^-2^ dyn.cm^-2^ respectively. Upon increasing the shear rate in the range of 10^-2^ dyn.cm^-2^, cells were found to become rounder with smaller nuclei.

## 4. Discussion

### Migratory behavior

Invasive tissue migration, *via* metastasis, is a mortality factor in cancer. After seeding on ECM models, ovarian tumor spheroids migrated isotropically, radially from the center of the spheroid. This is attributed to the chip geometry as well as the low shear stress imposed in our ovarian cancer-on-chip, in which smaller channels at the entrance to the central chamber spill the flow. In addition to being within the physiological range of the shear stress at play in the peritoneal cavity, isotropic migration is essential as it mimics the ascitic peritoneal environment, where the shear stress resulting from walking and breathing movements is effectively isotropic [3]. The migratory behavior was evaluated by measuring the distance of migration, the number of individual migrating cells out of the core of the spheroid and the core area. All of these descriptors were found to be strongly affected by the composition of the ECM model made up of collagen 1-based connective tissues (ACT) topped or not with a basement membrane from type IV and laminin (ABM). Because these two proteins are degraded in the course of ovarian cancer, these two ECMs reproduce the late and early stages of ovarian cancer progression respectively. An individual migration mode, characterized by a small area of the spheroid core and numerous individual migrating cells was observed on ACT, while a more collective behavior was observed for an early migration stage, on the ABM (Figure 9A). In this case, large spheroid cores with a limited number of individual cells were observed. This highlights the importance in considering the nature of ECM to elaborate models. Increased migration is often associated with an increase in stiffness in collagen or collagen-Matrigel mixture [30,31,47]. In our work, the stiffness of ACT and ABM, as measured by AFM nanoindentation, showed non-significant differences with mean values of 17 and 12 kPa for ACT and ABM respectively [32]. The difference in migratory behavior is rather attributed here to the cell-ECM interactions in presence of type IV collagen and laminin in the ABM model and the ability of this high-density coating of retaining factors from the culture medium as previously observed with Matrigel [30,31].

**Figure 8.**
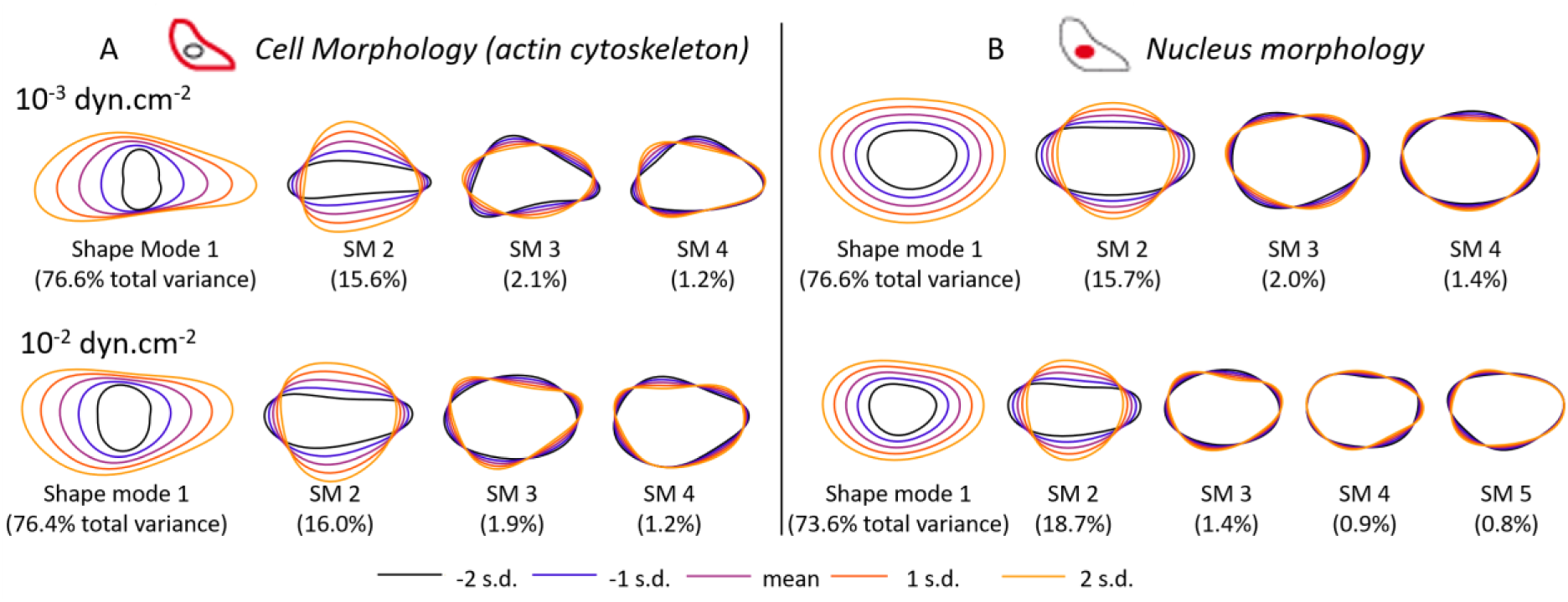
PCA of nucleus and cell morphology 48h after seeding on ABM as a function of flow shear stress: (A) morphological variance of cell morphology based on phalloidin staining of the actin cytoskeleton and (B) morphological variance of nucleus morphology based on DAPI staining. The color code is common to all plots. s.d. is for standard deviation, as determined by PCA for the quantification of the variance in the measured morphological features.

**Figure 9.**
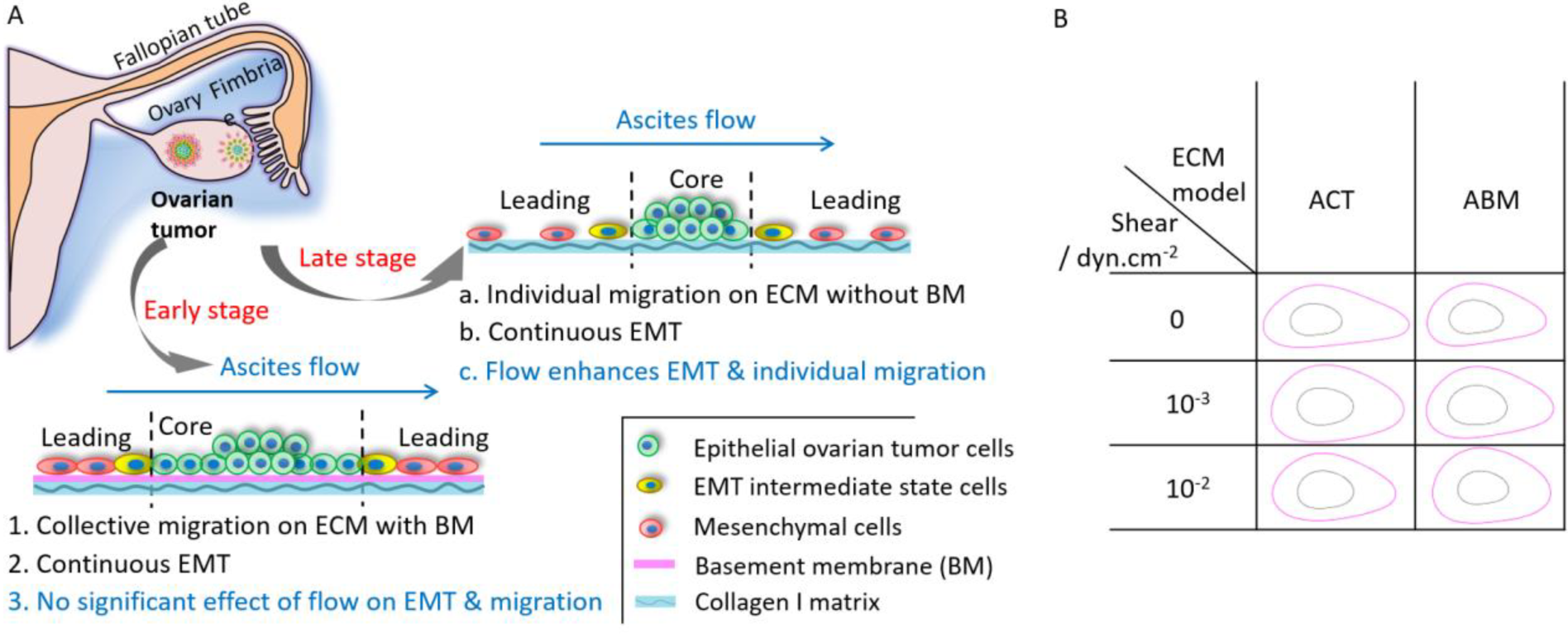
(A) Scheme of the effect of ECM remodeling and flow shear stress on the EMT and metastasis of OC cells. (B) Scheme of nucleus and cell morphology evolution of SKOV-3 cells on ACT and ABM in different shear stress conditions.

Interestingly, this strong difference in migration behavior depending on the ECM model was preserved under flow. Individual migration with decreasing area of spheroid core was observed on ACT with increasing flow rate, while a large core characteristic of a more collective migration mode was observed whatever the shear stress. This is consistent with the activation of integrins and focal adhesions in mechanotransduction pathways governing cell migration under flow shear stress [48]. This should account for the dominance of adhesion in regulating cell migration in the course of ECM remodeling during cancer progression, beyond its stiffening, with the loss of the basement membrane [49].

### Cytoskeleton reorganization and morphology

The intensity of vimentin staining was found to increase from the core to the leading area in all conditions, whatever the ECM model and shear stress applied. This illustrates the reorganization of the cytoskeleton of epithelial type of cells in the core of the spheroid transitioning towards mesenchymal type of cells with enhanced migratory properties. This is a continuous process at the origin of the cellular heterogeneity revealed by PCA analysis. Interestingly, flow shear stress enhances vimentin staining in SKOV-3 cells seeded on ACT, while a limited effect could be measured on ABM. In terms of morphology, a high dependency on ECM was observed in static conditions, cells being larger and more elongated on ACT than on ABM. Overall, in static conditions, cellular heterogeneity was found to be higher, with more shape modes recapitulating 95% of variations. Under flow, variations in cell morphologies were found to be lowered, and cells become smaller and rounder on both ECMs compared to the static conditions (Figure 9B). It is worth mentioning that from 10^-3^ to 10^-2^ dyn.cm^-2^, ACT cells preserve an elongated shape.

Overall, ECM plays a crucial role in determining whether ovarian cancer cells migrate individually or collectively. The addition of fluid-induced shear stress had a different impact on cell migration depending on the nature of the ECM. On ACT, shear stress enhanced individual migration, with a decrease in the number of migratory cells, a decrease in spheroid core area and an increase in the intensity of vimentin staining, while a limited impact of shear stress was observed on ABM collective migration. This is explained by the self-protecting features of collective assemblies, where cells in the spheroid core are protected by surrounding cells, themselves less exposed to shear due to enhanced cell-cell interactions. In contrast, individual cells are more exposed to shear and therefore more likely to be affected by an increase in shear stress.

The ability of our models not only to discriminate between ECM and flow shear stress, but also to investigate both factors synergistically is an important asset as flow shear stress has been studied with little attention to ECM evolution. While physiological 3D connective tissue models [48,50], including those using Matrigel [31] have been combined with flow shear stress, the use of a controlled number, concentration and organization of several ECM components (type I and type IV collagen, laminin) considerably refines the model, particularly for studying cancer at different stages. This is a key parameter to hold when designing therapeutically relevant models, given the reported impact of flow in therapeutic resistance in ovarian cancer [51].

## 5. Conclusion

We have built an easy-to-handle ovarian tumor-on-chip to reproduce the ECM environment and ascitic flow conditions during ovarian cancer. This device highlights the importance of key basement membrane proteins, type IV collagen and laminin, in the migratory behavior of cells. Indeed, ECM was found to prevail over shear stress in the physiological range of the peritoneal cavity in determining the individual or collective migratory behavior of ovarian cancer cells. Collective migration was observed on early-stage ECM, while a more individual migration was observed at later stage correlating with BM degradation. Furthermore, the application of shear stress did not have the same impact on migration depending on the ECM, with individual migration being enhanced by shear, while a limited impact was observed on more collective migrations. This work highlights the need to take ECM and its synergy with flow shear stress into account when studying cell migration, and even more so when developing tumor microenvironments-on-chip. The use of multiple ECM components has rarely been considered so far, and will be of paramount importance in the study of cancer stage progression for academic research and therapeutic developments.

## Credit author statement

## Acknowledgments

We thank the China Scholarship Council for the PhD grant of Changchong Chen; Rachele Allena and Nicolas Ruyssen for their help with shear stress simulations; Christophe Hélary and Gervaise Mosser for their help in the extraction and purification of type I collagen, and Clothilde Raoux for her constant support with SHG/2PEF experiments. This project has received financial support from the CNRS through the MITI interdisciplinary programs. Multiphoton imaging at LOB was partly supported by the Agence Nationale de la Recherche (contracts ANR-11-EQPX-0029 Morphoscope2 and ANR-10-INBS-04FBI). This work benefited from the technical contribution of the Institut Pierre-Gilles de Gennes joint service unit CNRS UAR 3750. The authors would like to thank the engineers of this unit for their advice during the development of the experiments: Bertrand Cinquin, Nhung Dinh, Audric Jan, Kévin Phan. We thank members of MEC-uP Team for helpful discussions and comments. This work was funded by ANR JCJC Modulo-EMT ANR-21-CE19-0006 and is a part of Alphonse Boché doctoral work.

