## Supplementary Information for "The balance between shear flow and extracellular matrix in ovarian cancer-on-chip"

### 1. Materials and methods

#### 1.1. Fabrication of the porous supports

#### 1.2. Spheroid growth

#### 1.3. Simulating the flow-induced shear stress of the peritoneal cavity

### 2. ABM characterization using two photon microscopy

### 3. E-cadherin and EpCAM staining of SKOV-3 cells

### 1. Materials and methods

#### 1.1. Fabrication of the porous supports

Porous supports similar to the ones that are commercially available from the Mesobiotech® company were fabricated by photolithography and soft-lithography in the lab as previously.[28] Supports were designed using CleWin5 software and printed from micropattern generator (Heidelberg µPG 101 Tabletop Micro Pattern Generator) with an array structure with a honeycomb microframe. The width of the honeycombs was set at 400 µm, with a frame width of 50 µm and a thickness of 50 µm. First, a double layer SU-8 mold was fabricated by photolithography. The mesh layer was patterned on a silicon wafer using a 50 µm thick SU-8 negative photoresist by UV exposure at 250 mJ/cm². Then, the honeycomb frame of 50 µm height was directly patterned on the mesh layer by another round of UV exposure at 250 mJ/cm². After development in propylene glycol methyl ether acetate, this double layer SU-8 mold was then exposed in trimethylchlorosilane (TMCS, Sigma, France) vapor for 10 min. Afterwards, a mixture of PDMS (GE RTV 615) pre-polymer and its crosslinker at ratio of 10:1 (w/w) was casted on the SU-8 mold. After curing at 75°C for 4 hours, the PDMS layer was peeled off. Then, this replicated PDMS structure was placed on a glass plate and a solution of a photo-crosslinking polymer (Ormostamp®, micro resist technology) was injected in the free space of the PDMS-glass assembly, followed by UV exposure at 1500 mJ/cm², and removal of the PDMS mold. The porous supports were then coated with gold by sputter deposition for improving hydrophilicity and drop casting of type I collagen by using an Emitech K675X Sputter Coater System working at 125 mA for 30 seconds.

#### 1.2. Spheroid growth

Ovarian tumor spheroids are produced using a caged-honeycomb supports with a non-adherent coating to control their growth. First, gelatin nanofibers and agarose are coated on one side of the porous scaffold with 200 µm thickness. Second, the coated porous scaffold is maintained by a PDMS ring at the bottom of a 12-well plate (Figure S1). Finally, the SKOV-3 cell suspension is cast into the hole of the PDMS ring with cell culture medium. SKOV-3 spheroids with a stable size are obtained after 3 days of culture.

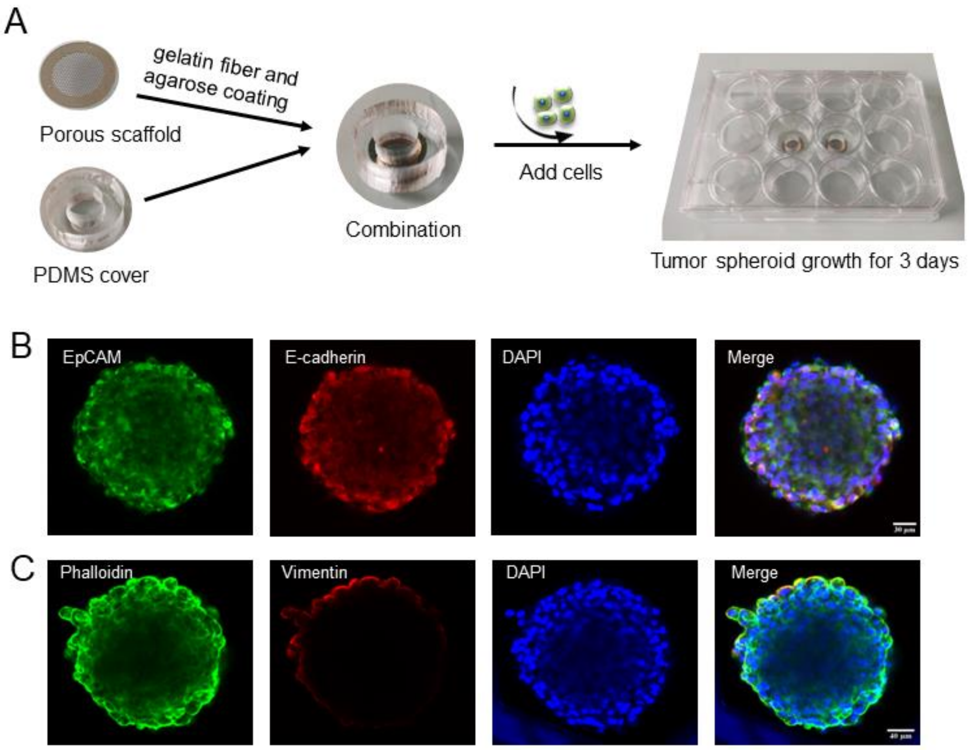


**Figure S1.** (A) Scheme of the preparation of spheroids. IF confocal imaging of a SKOV3 tumor sphere stained (B) epithelial (EpCAM and E-cadherin), and (C) mesenchymal markers (vimentin). The scale bar is 40 µm.

Figure S2 shows the resulting SKOV-3 spheroids with the uniform size of about 200 µm. The obtained spheroids can be extracted and seeded one by one. Immunofluorescence (IF) of the resulting spheroids confirms their 3D organization, as well as the presence of the epithelial markers EpCAM and E-cadherin. In contrast, staining of the vimentin cytoskeleton protein, which is commonly used as a mesenchymal marker could not be identified (Figure S2). After seeding, tumor spheroids were incubated on the ECM models for 2 hours before their transfer in the chip for culturing under perfusion for two days.

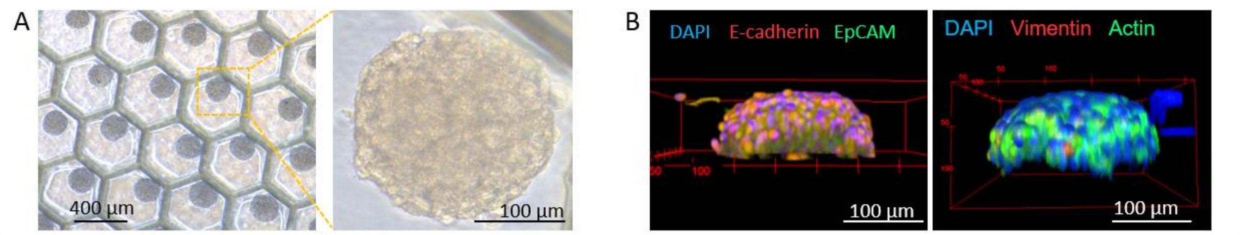


**Figure S2.** (A) Optical microscopy and (B) IF images of spheroids grown in deep-well patches. Cells are stained for DAPI, E-cadherin, EpCAM (left image), and for DAPI, vimentin and actin (right image).

#### 1.3. Simulating the flow-induced shear stress of the peritoneal cavity

Based on similar setups in the literature, we chose our flow to be around some tens of µL/min, as it matches with a physiologically relevant shear stress less than a $dyn.cm^{-2}$ for our setup. Besides, with our values, our flow was laminar according to (Eq.1).

$$Re = \frac{u_{(cm/s)} * L_{(cm)} * \rho_{(kg/m^3)}}{10 * \mu_{(dyn.s/cm^2)}} < 2000 \qquad \text{(Eq.1)}$$

where µ is the dynamic viscosity of water ($\mu \approx 0.01$ $dyn.s/cm^2$ at 20° C), L the typical dimension of the channel (L = 0.5 mm) and ρ the density of water (ρ = 998 $kg/m^3$ at 20° C). Consequently, computer fluid dynamics modeling for laminar flow was used. We assumed our fluid to be incompressible and Newtonian, and our system to be time independent. The structure of our chip was built with rectangular channels of width and height 500 µm, and central cavity of radius 4 mm and height 900 µm. The patch was superimposed to the central cavity and was 200 µm high. The inlet channel, which goes along the x axis (with z axis being defined orthogonally to the chip) is divided into six smaller channels to spill the flow into the central cavity while reducing the shear stress. As simulated by computer fluid dynamics modeling, this allows the shear stress due to the fluid to be homogenized within the chip. Because of the symmetry properties of our chip, only half of the chip was modeled to ease computations, with a condition of symmetry in the plan y = 0. Other boundaries were considered as no-slip walls, except from the inlet and outlet (the difference in pressure was defined as pressure imposed by the syringe actuator minus ambient pressure). The patch was approximated by a porous layer of collagen I, while the chip was made of PDMS, and filled with water. A single-phase laminar flow model was used to describe the flow, coupled with a Darcy's law in the porous area at the top of the central cavity. Porosity and viscosity required for the simulation were determined from material. A physics-controlled mesh with fine element size and without further modifications was used. Results were analyzed by plotting iso-pressure lines as well as velocity and shear stress mapping with different initial pressure (Figure S3).

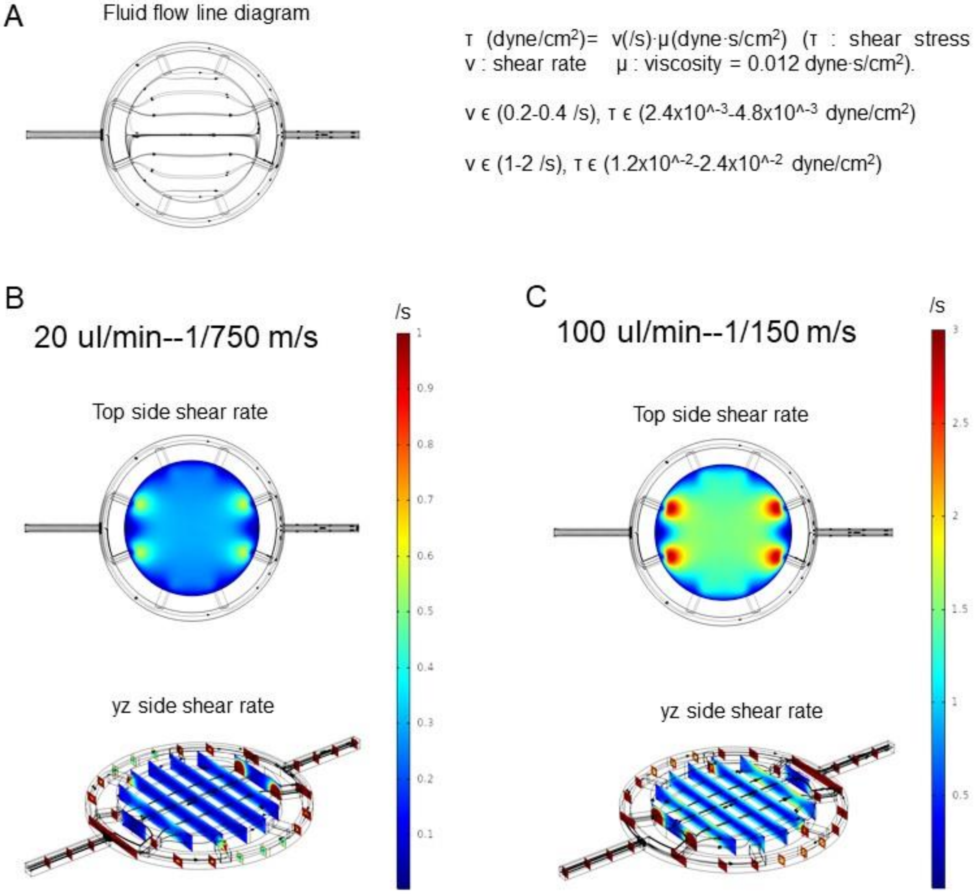


**Figure S3.** (A) Simulation of fluid flow direction and distribution in circular microchamber using Comsol Multiphysics. (B,C) Distribution of the top wall shear stress in the chamber for flow rates of 20 and 100 µl.min$^{-1}$.

### 2. ABM characterization using two photon microscopy

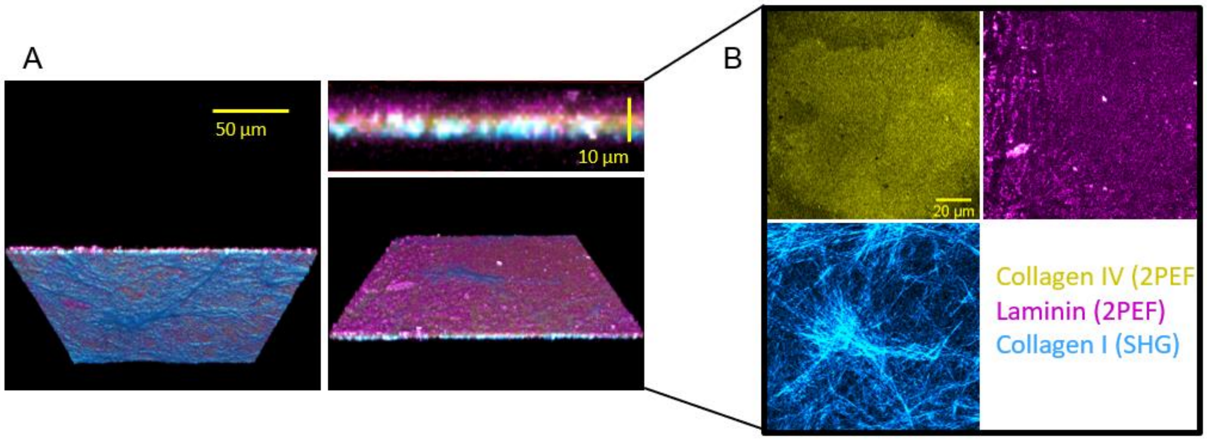


**Figure S4.** (A) SHG (collagen I in cyan) and 2PEF (laminin in magenta and collagen IV in yellow) images of the ABM. The left image is the bottom view, the top right one the side view, and the bottom right image is the top view. The underlying gelatin layer is not stained. (B) Details of the single multiphoton channels.

#### 3. E-cadherin and EpCAM staining of SKOV-3 cells

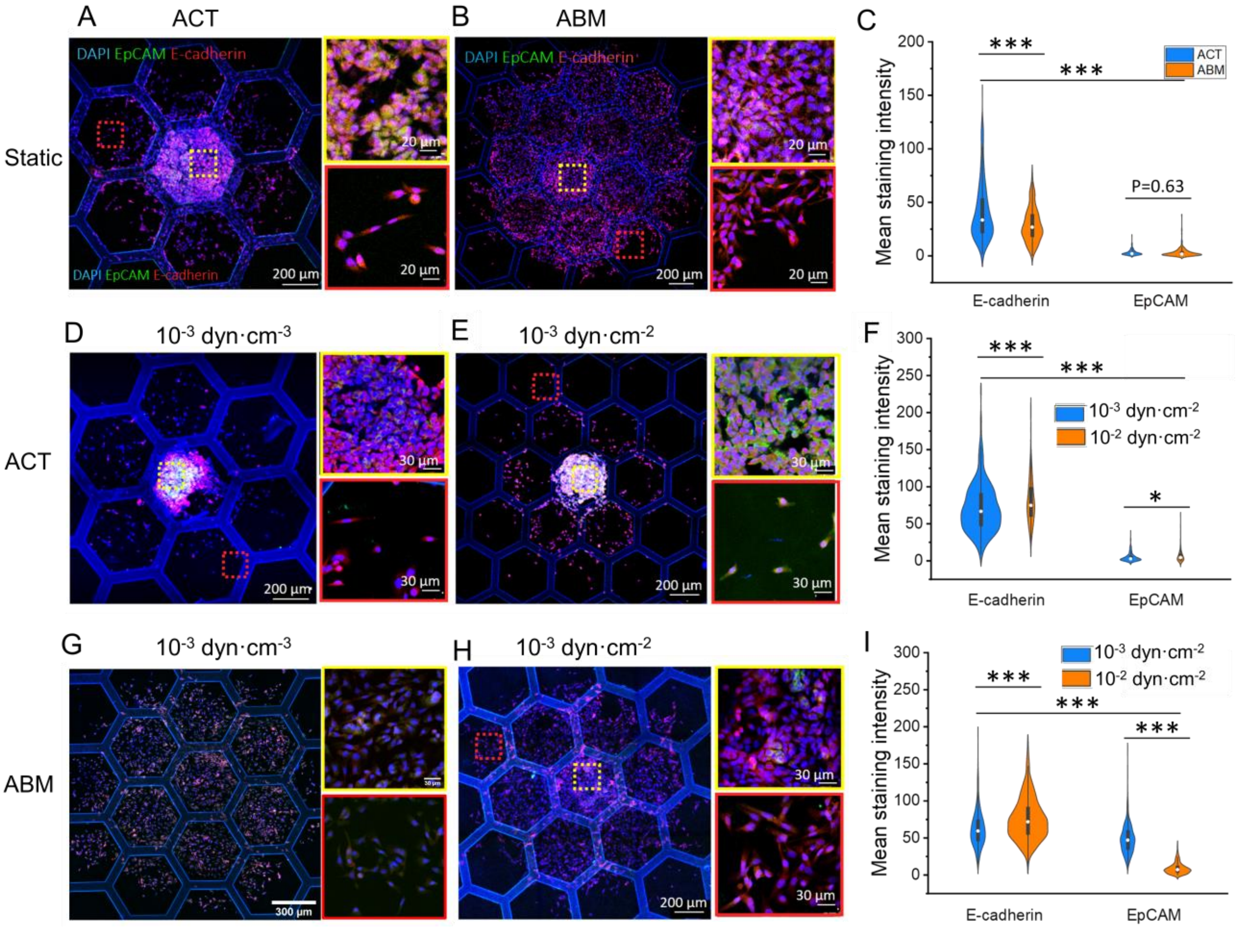


**Figure S5.** Effects of flow shear stress on migration of SKOV-3 cells on (A, D and E) ACT and (B, G and H) ABM. IF imaging of SKOV-3 cells after 48 hours culture on ABM. Cells were stained for E-cadherin (red), EpCAM (green), and nuclei (DAPI, blue). For epithelial markers, we used EpCAM CD326 Monoclonal Antibody (1B7) (Alexa Fluor 488, eBioscience™ Catalog # 53-9326-42) at a 1/20 dilution overnight at 4°C, and E-Cadherin (ab194982) at 1/250 dilution overnight at 4°C. C, F and I are the mean staining intensity of epithelial makers EpCAM and E-cadherin in different culture condition (* p<0.05, * * p<0.01, * * * p<0.001.)

The staining intensity of epithelial markers, E-cadherin and EpCAM, was found to decrease steadily from the core to the front at all shear stress. This supports the transition of SKOV-3 cells towards mesenchymal phenotypes from the spheroid core to the migration front, whatever the perfusion speed.
